# *TMEM230* is not a gene for Parkinson’s disease

**DOI:** 10.1101/097030

**Authors:** Matthew J. Farrer, Austen J Milnerwood, Jordan Follett, Ilaria Guella

## Abstract

Deng *et al.* report the discovery of *TMEM230* c.422G>T (p.Arg141Leu) mutation as a cause of late-onset, autosomal dominant Parkinson’s disease (PD)^1^ in the same pedigree in which we previously assigned *DNAJC13* c.2564A>G (p.Asn855Ser) as pathogenic^2^. The chromosome 20pter-p12 locus was discovered by short tandem-repeat (STR) genotyping and linkage analysis, with subsequent exome sequencing in four affected (II-4, III-1, III-20 and III-26) and one unaffected family member. Deng *et al.* state the rationale for their re-analysis was the inconsistency of genotype-phenotype correlations for *DNAJC13* c.2564A>G (p.Asn855Ser), this mutation being absent in three affected family members (II-1, III-1 and III-23). However, two of these suffer atypical parkinsonism; II-1 had clinical and pathologically-proven progressive supranuclear palsy, not Lewy body PD, whereas his son developed symptoms more than two decades earlier than the mean age at onset in the family. Based on haplotype and Sanger sequence analysis the authors’ claim *TMEM230* c.422G>T fully co-segregates with disease. Further discussion of these authors’ claims is necessary.

Our prior exome analysis of III-15 identified a *DNAJC13* c.2564A>G (p.Asn855Ser) mutation but not *TMEM230* c.422G>T (p.Arg141Leu) (Figure 1a). The exons of both genes had >30x sequence coverage and nucleotide sequences at both sites were confirmed by Sanger sequencing. In addition, our chromosome 20 STR marker genotyping, while consistent with parental/sibling haplotypes and Mendelian inheritance (Figure 1a) shows the genotyping presented by Deng *et al.* is erroneous for two individuals; III-14 at age 75 years is an unaffected carrier of TMEM230 c.422G>T (p.Arg141Leu) whereas III-15 is wild-type (Figure 1a, compare with Supplementary Fig. 1^1^). Many of the samples we have examined were obtained twice including III-15; blood was collected prior to 2010 and at a family reunion organized in 2013, and these samples gave identical results. Nevertheless, phenocopies in large pedigrees with multi-incident, monogenic late-onset parkinsonism are to be expected^3^. Deng *et al.* state *DNAJC13* c.2564A>G is present in an asymptomatic individual (II-9), who died at 87 years, and they excluded variants present in this exome and shared by the intersection of four affected exomes. We were not aware of a DNA sample from this subject but as reduced penetrance is a feature of monogenic parkinsonism^4^ we appreciate ‘disease-discordant’ analysis is best avoided. Indeed the authors’ observe *TMEM230* c.422G>T (p.Arg141Leu) in 20/23 subjects including 12 unaffected carriers in generation III (Supplementary Fig. 1^1^, and correcting for the genotyping error in III-14; Table). The number of carriers is consistent albeit greater than one might expect for a dominant pattern of disease inheritance.

**Figure 1a.**
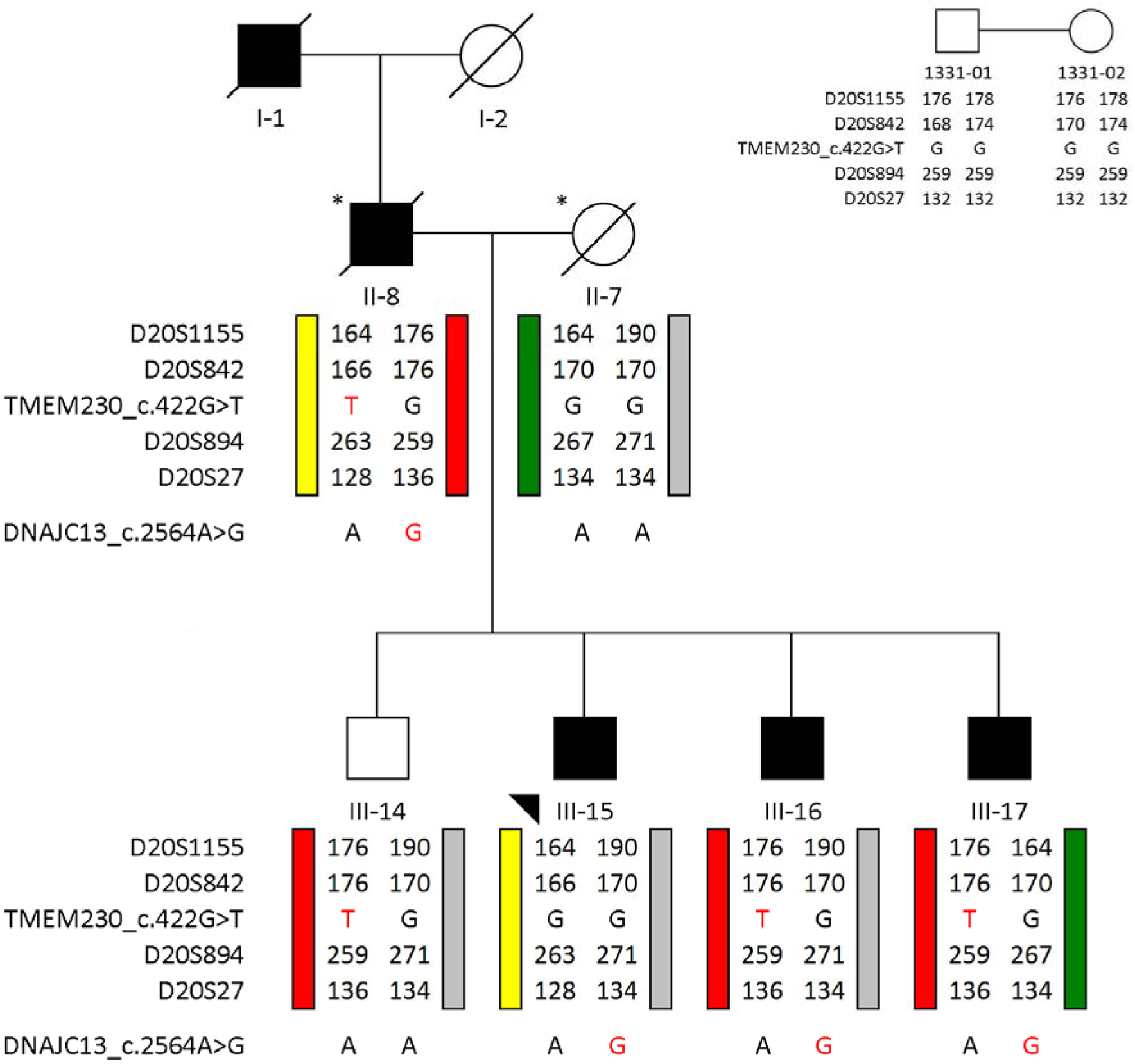
Chromosome 20pter-p12 genotyping and mutation analyses. Chr20pter-p12 haplotypes are color-coded according to Deng *et al.* Genotype results for the locus include four STR markers with allele sizes according to CEPH standards (inset, right). *TMEM230* c.422G>T (p.Arg141Leu) and *DNAJC13* c.2564A>G (p.Asn855Ser) mutations are displayed. Affected individuals are shown as filled black symbols. The arrowhead indicates the exome-sequenced individual. DNA samples are available on request, or directly from Dr. Ali Rajput, Royal Hospital, University of Saskatchewan. An asterisk indicates DNA samples for whom genotypes are inferred.

**Table.**
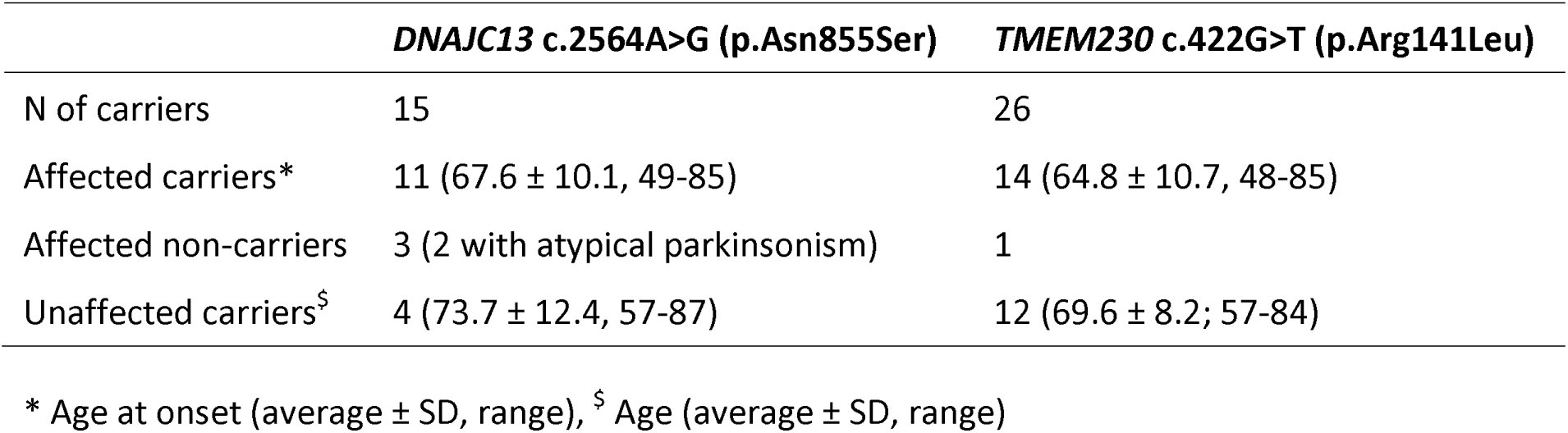

|  | ***DNAJC13* c.2564A>G (p.Asn855Ser)** | ***TMEM230* c.422G>T (p.Arg141Leu)** |
| --- | --- | --- |
| N of carriers | 15 | 26 |
| Affected carriers* | 11 (67.6 ± 10.1, 49-85) | 14 (64.8 ± 10.7, 48-85) |
| Affected non-carriers | 3 (2 with atypical parkinsonism) | 1 |
| Unaffected carriers<sup>§</sup> | 4 (73.7 ± 12.4, 57-87) | 12 (69.6 ± 8.2; 57-84) |

\* Age at onset (average ± SD, range), <sup>§</sup> Age (average ± SD, range)

However, the authors’ fail to disclose that many *TMEM230* c.422G>T carriers are advanced in age with 7 individuals beyond the mean age of symptom onset (mean age of unaffected carriers = 69.6±8.2, range 57-84 years; Table). Given these issues segregation of the *TMEM230* c.422G>T (p.Arg141Leu) with parkinsonism is arguably less compelling than for *DNAJC13* c.2564A>G (p.Asn855Ser). Two-point linkage analysis of the chromosome 3q21.3-q22.2 disease-segregating haplotype (also harboring and *ZBTB38* c.791G>A (p.Arg264Gln) variants, 9Mb apart) yielded a LOD=4.79 (θ=0.05) which increased to 5.29 (θ=0) assuming a 10% phenocopy rate. Hence, other considerations for the pathogenicity of these variants must be carefully reviewed.

The *DNAJC13* c.2564A>G mutation is not observed in public databases (ExAC 0.3.1 release Mar 14th 2016) and the amino acid codon it encodes, p.Asn855, is evolutionarily conserved across species suggesting it is important to protein function. In contrast, while *TMEM230* c.422G>T is novel its codon p.Arg141 is not evolutionarily conserved in humans. This is an important caveat. TMEM230 p.Arg141Gln and p.Arg141Trp substitutions occur with appreciable frequency; in Caucasians their combined minor allele frequency (MAF) > 5e^-5^ and in South Asia TMEM p.Arg141Trp MAF=0.00024 alone (ExAC 0.3.1 release Mar 14th 2016). CADDv13^5^, which provides a composite prediction of damage to protein structure, suggests these substitutions are comparable (CADDv13 scores for DNAJC13 p.Asn855Ser =23.6, and for TMEM230 p.Arg141Leu, p.Arg141Gln and p.Arg141Trp = 27.6, 23.8 and 28.6 respectively). While the MAF for TMEM p.Arg141 substitutions is more than one might envisage for a rare cause of parkinsonism the clinical phenotype of these carriers is unavailable and *in-silico* predictions cannot be assumed causal of disease. Thus Deng *et al.* may find it informative to compare the consequence of these and more frequent TMEM230 substitutions in their functional assay (see below).

For Nature Genetics ‘proof of pathogenicity’ requires additional mutations in the same gene segregate with disease in independent families. In our original study screening of *DNAJC13* c.2564A>G (p.Asn855Ser) identified five more affected carriers but no controls; three of the positive cases have a family history of parkinsonism and in two small pedigrees co-segregation with PD was evident. All DNAJC13 p.Asn855Ser carriers are of Dutch-German-Russian Mennonite origin and share a common haplotype^2^. Our screening of *TMEM230* c.422G>T in 1283 Canadian patients (726 with sporadic PD and 557 with familial parkinsonism), including those of Dutch-German-Russian Mennonite ancestry in our original report, did not identify any carriers. Deng *et al.* go on to identify *TMEM230* c.275A>G (p.Tyr92Cys) and c.551A>G (p.*184Trpext*5) in two patients with early-onset parkinsonism; the former observed in an unaffected maternal carrier and the latter found in a sporadic case. In addition, *TMEM230* c.550_552delTAGinsCCCGGG (p.*184ProGlyext*5) was found in affected probands from 7 of 225 unrelated families of ethnic Han Chinese descent. Nevertheless, while all these mutations are novel, none are shown to segregate with disease. TheTMEM230 p.*184ProGlyext*5 discovery is most incongruous given the independent/ancient haplotypes on which it is observed (Supp. Fig 12d) and the 3.1% of familial PD in ethnic Chinese it is stated to explain. In contrast to TMEM230 p.Arg141Leu the p.*184ProGlyext*5 substitution does not appear to follow a Mendelian pattern of inheritance, although the lack of clinical information for parents and siblings (many of whom must be carriers) offers little insight. We have examined *TMEM230* variability in 256 patients with early-onset, familial and/or atypical parkinsonism; while we observe 4/7 *TMEM230* variants leading to amino acid substitutions (p.Met1Val, p.Arg62His, p.Ile125Met, p.Arg171Cys) all have appreciable frequencies in ExAC (MAF >5e^-5^) and only p.Arg171Cys is predicted to be damaging to protein structure (MAF=0.003, CADDv13 score=34). We also genotyped *TMEM230* c.550_552delTAGinsCCCGGG in 80 affected probands with familial PD and 280 control subjects of ethnic Chinese descent from Taiwan but found no carriers.

Deng *et al.* claim TMEM230 mutations impair ‘synaptic vesicle trafficking’^1^. However, assuming the TMEM230 antibodies are specific (which was not demonstrated in the manuscript), TMEM230 localized to large vesicular structures in the cell body. The localization appeared similar for endogenous and overexpressed/tagged protein in Neuro-2a cells and brain tissue (Figures 2 &3^1^). Similarly, confocal live cell imaging showed the movement of tagged, overexpressed (non-physiological) proteins in neuronal soma (Figure 4^1^). Assessing ‘synaptic vesicle’ transport speed, track length and displacement length is claimed to show some mutant specific differences. Nevertheless, synaptic vesicles by definition reside in presynaptic axon terminals; somatic membrane structures containing vesicular proteins are not synaptic vesicles which form from membrane intermediates at axonal sites hugely distal from the soma. It is incongruous this biology was not studied in an appropriate, physiological context or at least termed correctly. How images were quantified is unclear; at a minimum transfection levels for each tagged-protein must be comparable and should have been demonstrated. The significance of the 1-way ANOVA is also based on comparison to an overexpressed, tagged wild-type protein. The more frequent and presumably non-pathogenic TMEM230 substitutions we have identified might prove to be informative as controls.

In contrast, we have studied the consequences of the *DNAJC13* c.2564A> G (p.Asn855Ser) mutation in synaptically mature primary cortical cultures (DIV21) prepared from a knock-in mouse (Figure 1b). We observe profound effects on endosomal tubulation in heterozygotes, and in endosomal proliferation in both heterozygous and homozygous mutant animals, with respect to neurons grown from wild-type embryos from the same litters. Our findings recapitulate and extend observations made in DNAJC13/RME-8 knockdown experiments in immortalized cell lines using the same SNX1-GFP endosome marker^6^ suggesting a gene-dose dependent ‘loss of function’ is induced by this mutation.

**Figure 1b.**
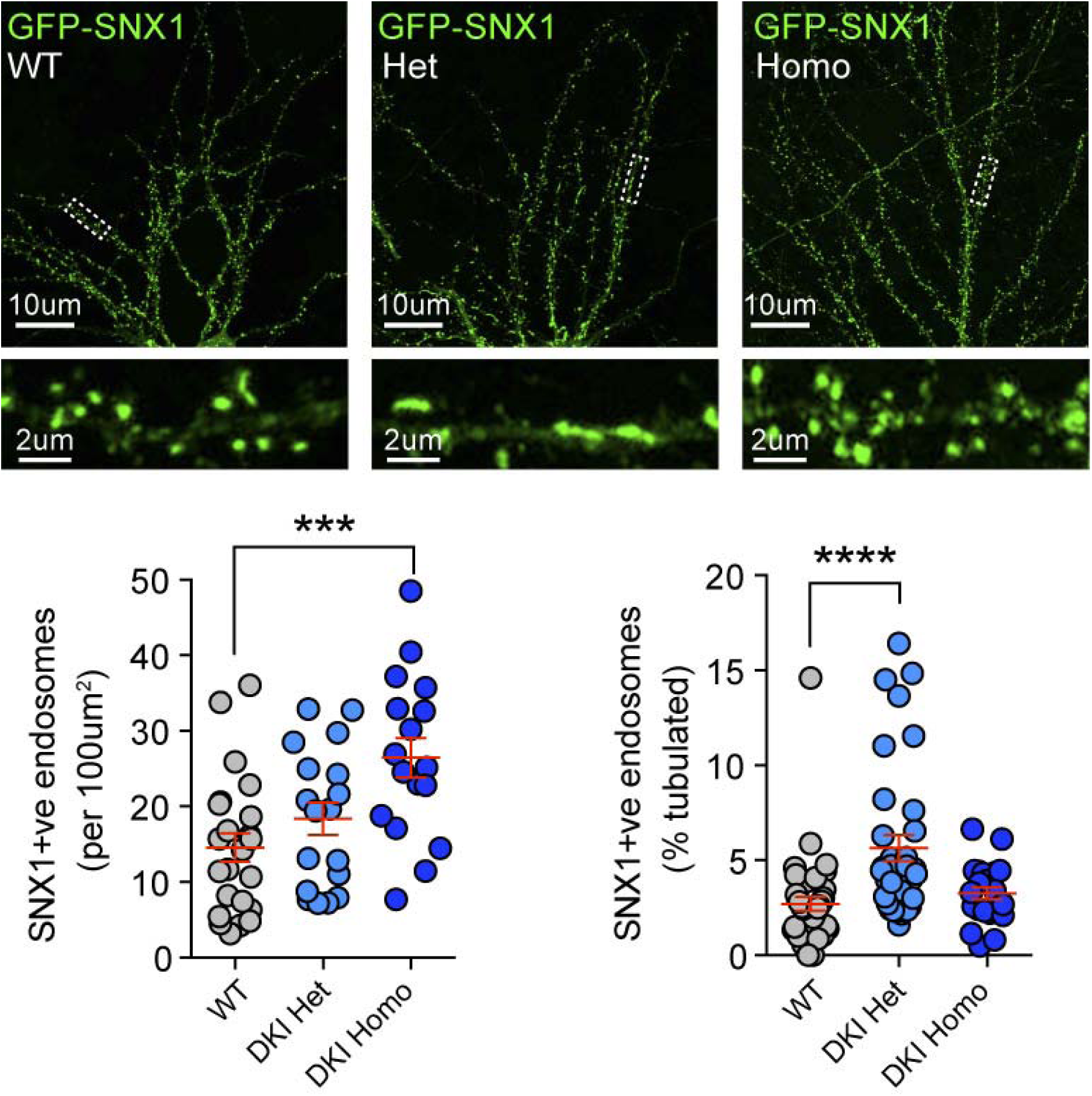
Impaired endosomal tubulation in DNAJC13/RME-8 p.Asn855Ser knock-in (DKI) mice. DNAJC13/RME-8 p.Asn855Sr knock-in (DKI) mice were engineered on a C57BL/6JN background by Flp-deleter transgenesis (Taconic). Brain tissue was collected as previously^7^ from 3 month WT and DKI littermates, flash frozen, and lysed in HEPES buffer (20mM HEPES pH7.4, 50mM CH_3_C0_2_K, 200mM Sorbitol, 2mM EDTAand 0.1% Triton X-100) with protease inhibitors (Roche). Lysates were denatured (lamelli buffer 95C for 5 minutes), run on NuPage Bis-Tris gel, and transferred to Immobilon FL membranes. Membranes were probed with validated rabbit anti-RME-8^2^ and mouse anti-GAPDH (Thermo Scientific) primaries followed by 488-conjugated secondaries and imaged (Chemidoc MP, BioRad). Band intensities were measured in ImageLab and RME-8 signal normalized to GAPDH in each lane. Cortical primary neuronal cultures were prepared as previously^8^ from single wild-type littermate (WT) and DKI pups. Cells were pooled by genotype and nucleofected with plasmids to express the endosome marker GFP-SNX1 under a synthetic CAG (CMV/chicken ß-actin) promoter (Addgene 37825) prior to plating, and maintained to 21 days in vitro (DIV21). At DIV21 cells were fixed, stained with anti-GFP (rabbit) primary and Alexa_48_ secondary antibodies, slide mounted and imaged by confocal laserscanning microscopy (Olympus Fluoview1OOO, 60x, 2x zoom, 1um z-stacks). Stacks were flattened, background subtracted and binarized prior to particle analysis (10 pixel rolling ball subtraction algorithm, ImageJ version 1.15d) to determine density, size and % of tubulated (<0.5 circularity) GFP-SNX1 positive endosomes in masked primary, secondary, and tertiary dendrites. Data was collated and statistical analyses performed by 1-Way ANOVA and significant differences reported from Dunn’s (% tabulated) and Dunnett’s (density) post-tests in Graphpad Prism6.

In conclusion, *DNAJC13* c.2564A> G (p.Asn855Ser) remains compelling as the gene mutation for familial parkinsonism in this Mennonite kindred. Future research in DNAJC13 biology will undoubtedly make important contributions to the field, for which DNA constructs and the constitutive knock in mouse model are now available (on request). In contrast, the evidence *TMEM230* is a gene for familial parkinsonism is not compelling. The mutations Deng and colleagues describe fail to segregate with disease, as claimed. TMEM230 p.Arg141Leu, at non-conserved amino acid residue, is also rather too frequent to be causal of disease. Neither is there any evidence to support the notion that *DNAJC13* and *TMEM230* mutations may act together, in concert, to lead to parkinsonism. We recommend Deng and colleagues reassess and retract their prior work. In the interim, we invite independent authors to take the opportunity to assess our genetic analysis and samples (Figure 1a), and publish their findings accordingly.

## Acknowledgements

Original data is contributed with technical support from Adam Book, Stephanie Bortnick, Liping (Daisy) Cao, Dan Evans, Chelsie Kadigen and Lise Munsie on behalf of the Centre for Applied Neurogenetics. We sincerely appreciate clinical samples and contributions by Drs. Alex and Ali Rajput (Royal Hospital, University of Saskatchewan, Saskatoon, Canada), Prof. Ruey-Mei Wu (Taiwan National University Hospital, Taiwan) and Drs. Silke Appel-Cresswell, Martin McKeown and Prof. A. Jon Stoessl (Pacific Parkinson’s Research Centre, University of British Columbia, Vancouver, Canada). We also thank Dr. Carles Vilarino-Guell for helpful discussion.

## References

1. Deng, H.X. et al. Identification of TMEM230 mutations in familial Parkinson’s disease. Nature genetics 48, 733–9 (2016).

2. Vilarino-Guell, C. et al. DNAJC13 mutations in Parkinson disease. Human molecular genetics 23, 1794–801 (2014).

3. Klein, C., Chuang, R., Marras, C. & Lang, A.E. The curious case of phenocopies in families with genetic Parkinson’s disease. Movement disorders: official journal of the Movement Disorder Society 26, 1793–802 (2011).

4. Trinh, J., Guella, I. & Farrer, M.J. Disease penetrance of late-onset parkinsonism: a meta-analysis. JAMA neurology 71, 1535–9 (2014).

5. Kircher, M. et al. A general framework for estimating the relative pathogenicity of human genetic variants. Nature genetics 46, 310–5 (2014).

6. Freeman, C.L., Hesketh, G. & Seaman, M.N. RME-8 coordinates the activity of the WASH complex with the function of the retromer SNX dimer to control endosomal tubulation. Journal of cell science 127, 2053–70 (2014).

7. Beccano-Kelly, D.A. et al. LRRK2 overexpression alters glutamatergic presynaptic plasticity, striatal dopamine tone, postsynaptic signal transduction, motor activity and memory. Human molecular genetics 24, 1336–49 (2015).

8. Beccano-Kelly, D.A. et al. Synaptic function is modulated by LRRK2 and glutamate release is increased in cortical neurons of G2019S LRRK2 knock-in mice. Frontiers in cellular neuroscience 8, 301 (2014).

